# Microbiome response in an urban river system is dominated by seasonality over wastewater treatment upgrades

**DOI:** 10.1101/2022.06.30.498375

**Authors:** Sho M. Kodera, Anukriti Sharma, Cameron Martino, Melissa Dsouza, Mark Grippo, Holly L. Lutz, Rob Knight, Jack A. Gilbert, Cristina Negri, Sarah M. Allard

## Abstract

Microorganisms such as coliform-forming bacteria are commonly used to assess freshwater quality for drinking and recreational use. However, such organisms do not exist in isolation; they exist within the context of dynamic, interactive microbial communities which vary through space and time. Elucidating spatiotemporal microbial dynamics is imperative for discriminating robust community changes from ephemeral ecological trends, and for improving our overall understanding of the relationship between microbial communities and ecosystem health. We conducted a seven-year (2013-2019) microbial time-series investigation in the Chicago Area Waterways (CAWS): an urban river system which, in 2016, experienced substantial upgrades to disinfection processes at two wastewater reclamation plants (WRPs) that discharge into the CAWS and improved stormwater capture, to improve river water quality and reduce flooding. Using culture-independent and culture-dependent approaches, we compared CAWS microbial ecology before and after the intervention. Examinations of time-resolved beta distances between WRP-adjacent sites showed that community similarity measures were often consistent with the spatial orientation of site locations to one another and to the WRP outfalls. Fecal coliform results suggested that upgrades reduced coliform-associated bacteria in the effluent and the downstream river community. However, examinations of whole community changes through time suggest that the upgrades did little to affect overall riverine community dynamics, which instead were overwhelmingly driven by yearly patterns consistent with seasonality. Such results emphasize the dynamic nature of microbiomes in open environmental systems such as the CAWS, but also suggest that the seasonal oscillations remain consistent even when perturbed.

**Importance:** This study presents a systematic effort to combine 16S rRNA gene amplicon sequencing with traditional culture-based methods to evaluate the influence of treatment innovations and systems upgrades on the microbiome of the Chicago Area Waterway System, representing the longest and most comprehensive characterization of the microbiome of an urban waterway yet attempted. We found that the systems upgrades were successful in improving specific water quality measures immediately downstream of wastewater outflows. Additionally, we found that the implementation of the water quality improvement measures to the river system did not disrupt the overall dynamics of the downstream microbial community, which remained heavily influenced by seasonal trends.

## Introduction

High quality fresh water is a critical natural asset that is under increasing risk of overuse and contamination by anthropogenic influences (Vörösmarty et al. 2010). With over half of the world’s total population living in urban areas, urban waterways are particularly influenced by human activity and act as a liaison between humans and the natural environment (Fisher et al. 2015). Historically, microorganisms have been utilized as metrics of water quality due to their traceability to potential sources of contamination. Concentrations of *Escherichia coli* or, more broadly, coliform-forming bacteria, have typically been used as indicators for fecal contamination in recreational, agricultural, and drinking water (McLain and Williams 2012). Although valuable, such approaches can vary in both reliability and accurate representation of human pathogen levels (Cabral 2010, Noble et al. 2003, McLain et al. 2011, Chung et al. 2020). Furthermore, culture-based methods do not capture changes in the ecological structure of the system, which can provide valuable insights into ecosystem health (Dorevitch et al. 2017, Rijal et al. 2009, Rijal et al. 2011). Bacterial pathogens and fecal indicator taxa do not exist in isolation; rather, they exist within expansive, interactive communities of diverse and abundant microbial members (Kodera et al. 2022). Therefore, improving our understanding of the relationship between microbiology and water quality may require a more holistic examination of the microbial community.

Microbial communities in aquatic systems can be highly responsive to environmental variables such as temperature, nutrient availability, hydrology, metal contamination, and contrasting land-use (Weller et al. 2022, Sungawa et al. 2015, Vasemägi et al. 2017, Zeglin 2015, Smith 2007, Singer et al. 2010). Water quality management efforts must therefore take into account such spatio-temporal dependencies, particularly in fluctuating environments such as riverine systems. To this end, we conducted a 7-year investigation (2013-2019) of the microbial community of the Chicago Area Waterway System (CAWS) (Figure S1a). The CAWS is an extensive, highly engineered urban river system running through Chicago that consists of over 76 miles of man-made canals and modified natural streams (Supplementary Figure S1). In collaboration with Chicago’s Metropolitan Water Reclamation District (MWRD) we collected monthly samples of water and sediment across 12 sites, as well as untreated sewage and treated effluent samples from two wastewater reclamation plants (WRPs). We analyzed these samples with 16S rRNA amplicon sequencing and fecal coliform enumeration, and continually monitored relevant physicochemical characteristics at each site. Notably, the sampling period of our study coincided with the implementation of two major water quality improvement efforts at the CAWS. In 2016, advanced disinfection systems at the two existing WRPs were implemented to improve treatment of wastewater before its discharge to the CAWS. A seven-channel UV chamber system was installed at O’Brien WRP, while a chlorination-dechlorination system was installed at Calumet WRP. Simultaneously, the segment of the Tunnel And Reservoir Plan (TARP) associated with the Calumet WRP was implemented (Figure S1b). The TARP is a tunnel system that captures combined untreated sewage and stormwater from the surrounding area. Without this intervention, Combined Sewer Overflow (CSO) directly discharges into the CAWS during rainy weather and high flow conditions. The implementation of these initiatives during our longitudinal study provided a unique opportunity to examine how microbial community dynamics in a wastewater-impacted water system may be affected when presented with substantial wastewater management upgrades.

The primary goals of this study were to a) characterize the microbial communities of the CAWS and its spatio-temporal dynamics across 12 riverine sites and 2 wastewater treatment plants for 63 timepoints over 7 years, b) compare such characterizations to the trends found in a traditional, culture-dependent approach to measuring water quality, and c) examine the impact of water quality improvement interventions on the microbial ecology of the CAWS. The combination of sequencing data, fecal coliform data, and physicochemical data thus enabled us to parameterize the microbial ecology of the CAWS to gain nuanced insights into how environmental disturbance impacts the ecology of this already dynamic microbial system.

## Materials and Methods

We utilized 16S rRNA gene amplicon sequencing, fecal coliform counts, and in-situ physicochemical measurements to characterize the microbial ecology of the CAWS from 2013-2019. Water and sediment samples were collected from 12 sites along the CAWS, while sewage and effluent samples were collected from the O’Brien and Calumet WRPs (Figure S1). We collected a total of 2,306 samples: 260 effluent samples, 558 sediment samples, 928 water column samples, 88 sewage samples, and 472 technical controls (bottle, filter, equipment blanks).

### Sample collection

All CAWS locations were grab-sampled monthly for water (*i.e*. river, effluent, sewage) and sediment by MWRD personnel. A total of 500 mL of surface water and 100 g of sediment were collected in sterile containers for microbiome analysis. Temperature, pH, conductivity, and turbidity of water samples were measured using a handheld YSI multiparameter digital water quality meter. NO_2_^-^/NO_3_^-^ ratios, NH_3_, and PO_4_^3-^ values were measured using a Lachat Quickchem 8500 Series 2.0 instrument, using EPA 353.2 Rev 2.0, EPA 350.1 Rev 2.0, and EPA 365.4 reference methods, respectively. Chlorophyll measurement protocols were adapted from standard methods used in the Examination of Water and Wastewater method 10200 H. Between 2013-2017 measurements were made using a Beckman DU-640 spectrophotometer, and in 2017-2019 measurements were made using a Thermoscientific Genesys 10s UV-VIS spectrophotometer. Between 2013 – 2018 dissolved oxygen was measured using the Winkler method for a fixed DO sample. After 2018, the measurement protocol was transitioned to using a HACH HQd portable meter with a luminescent DO probe. Sediment samples were stored in polypropylene containers at 4°C. Water samples (200 mL) were filtered in duplicate using 0.22 Micron Mixed Cellulose Ester filters, and filters were aseptically transferred to labeled sterile 50 mL tubes and stored at −80°C. All water and sediment samples were transferred to the lab on ice and then stored at −80°C until thawing for processing.

### 16S rRNA amplicon sequencing and bioinformatic processing

DNA was extracted using the protocol described by Marotz et al. (2017) and the V4 region of the 16S rRNA gene was amplified using the protocol described by Caporaso et al. (2012). Briefly, we used region-specific primers (515F-806R) that included the Illumina flow cell adapter sequences and a 12-base barcode sequence for amplification of each 25μl PCR reaction containing the following mixture: 12μl of MoBio PCR Water (Certified DNA-Free; MoBio, Carlsbad, USA), 10μl of 5-Prime HotMasterMix (1×), 1μl of forward primer (5μM concentration, 200pM final), 1μl of Golay Barcode Tagged Reverse Primer (5μM concentration, 200pM final), and 1μl of template DNA. The conditions for PCR were as follows: 94°C for 3 min to denature the DNA, with 35 cycles at 94°C for 45 s, 50°C for 60 s, and 72°C for 90 s, with a final extension of 10 min at 72°C to ensure complete amplification. Amplicons were quantified using PicoGreen (Invitrogen) assays on a plate reader, followed by clean up using the UltraClean^®^ PCR Clean-Up Kit (MoBio, Carlsbad, USA) and quantification using Qubit readings (Invitrogen, Grand Island, USA). Amplicons were sequenced on an Illumina MiSeq HiSeq2500 platform with paired-end sequencing at the Argonne National Laboratory Core Sequencing Facility according to protocols from the Earth Microbiome Project (Thompson et al. 2017).

The raw sequence data was demultiplexed, trimmed, and processed using the open-source microbial study management platform Qiita (Gonzalez et al. 2018). Parameters for quality filtering included 75% consecutive high-quality base calls, a maximum of three low-quality consecutive base calls, zero ambiguous bases, and minimum Phred quality score of 3 as suggested previously (Bokulich et al. 2013). Demultiplexed sequences were trimmed to 150 base pairs, and then selected for ASV (Amplicon Sequence Variant) picking using the Deblur pipeline v. 1.1.0 (Amir et al. 2017). In the pipeline, *de novo* chimeras were identified and removed, artifacts (i.e. PhiX) were removed, and ASVs in less than 10 samples were removed for further analyses due to low representation. Blank samples were used as negative controls to determine the read count threshold for downstream analyses. ASV tables were rarefied to a sequencing depth of 1285 reads, leading to the removal of 15 of 1834 non-blank samples which contained fewer reads than the set depth. Analysis was completed using both QIIME 2 2021.2 (Bolyen et al. 2019) and in R 3.4.2 via the phyloseq package (McMurdie and Holmes 2013).

### Alpha and beta diversity analysis

Alpha and beta diversity were calculated between different sample types as well as by year. Alpha diversity for all sample types was measured using Shannon’s index. Beta diversity was determined using both unweighted and weighted UniFrac distances (Lozupone and Knight 2005, Lozupone et al. 2007), which were ordinated using Principal Coordinate Analysis (PCoA). Statistical significance of the differences in microbial alpha diversity and beta diversity were assessed using paired t-tests with Benjamini-Hochberg corrections and permutational multivariate analysis of variance (PERMANOVA), respectively (Anderson 2017). Beta dispersion values for each sample type were calculated as the mean value of unweighted UniFrac distances between all coordinates of individual samples to the coordinate of its respective sample type centroid, with statistical significance assessed using paired t-tests with Benjamini-Hochberg corrections. Figures were generated using ggplot2() (https://github.com/tidyverse/ggplot2) in the R language (https://www.r-project.org/).

### Time series analyses

For each unique combination of sample type and site, the microbial community of a sample in early spring was designated as the “baseline” community for the sample type/site grouping (for effluent, water, and sediment, the baseline communities were set in March 2014; for sewage, the baseline communities were set in March 2015, as collection began after March 2014.). Next, the communities of all samples belonging to the same sample type/site grouping were compared to the baseline community through time using unweighted and weighted UniFrac distances. Distances to the baseline community (henceforth referred to as beta distances) were plotted through time to generate time series visualizations. Because no collections were conducted in the months of December, January, and February, missing time points were interpolated using linear spline interpolation (Moritz et al. 2015). Each time series plot was then decomposed into three components (trend, seasonality, and error) using STL (Seasonal-Trend decomposition procedure based on Loess; Cleveland et al. 1990). Analyses involving the identification of seasonal dynamics within sample types were then detrended according to the STL outputs (to remove any confounding effects of year on year changes in community dynamics), and the resulting time series were statistically tested for seasonality using Friedman tests with Benjamini-Hochberg corrections. Analyses and visualizations were performed using the tsutils (https://github.com/trnnick/tsutils) and forecast (Hyndman et al. 2022) packages in R.

### Differential abundance analysis

To identify specific microbial ASVs driving the observed patterns of seasonality in water and effluent samples while accounting for compositionality, we conducted differential abundance analyses on the non-rarefied dataset using Songbird v. 1.0.4 through QIIME2 v. 2021.2 (Morton et al. 2019). The differential abundance models were trained with the water and effluent samples using only the data from August and March time points. Songbird differential rankings were visualized in Qurro (Fedarko et al. 2020) to identify the 10 most differentially abundant ASVs in March relative to August, and the 10 most differentially abundant ASVs in August relative to March, separately with both water and effluent samples. Once March-associated and August-associated ASVs were identified, we examined changes in their log-ratios across the entire rarefied dataset of water and effluent samples to test whether the identified ASVs follow a predictable seasonal gradient through time. We used the following formula:

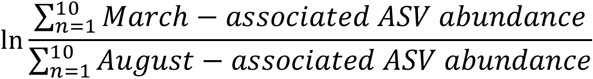

with a pseudo count of one applied to the table before taking the ratio to avoid undefined values.

### Statistical analysis of fecal coliform data

Due to the skewed sampling distributions and the presence of substantial outliers in the measured fecal coliform data, we employed a non-parametric, median-based bootstrap hypothesis test (Efron and Gong 1983, Manly 2007) to examine whether fecal coliform concentrations at select sites significantly changed post-intervention. At each site, the median fecal concentration value from time points post-intervention were contrasted with the median fecal concentration value from time points pre-intervention to generate a test statistic. Next, the fecal coliform data were randomly resampled with replacement among all the time points to generate a simulated test statistic given the null hypothesis that fecal concentrations are non-associated with the implementation of the intervention in 2016. This process was repeated 10,000 times to generate a null distribution of test statistics, which could then be compared against the true test statistic in a two-tailed test for significance. This analysis was performed with effluent samples, water samples immediately downstream from WRPs, and water samples further downstream from the WRPs.

### Statistical analysis of physicochemical data

We examined the effects of the wastewater disinfection treatments on the abiotic water characteristics of sites immediately downstream of the treatment plants. This was accomplished by testing for significant differences in key physicochemical variables (temperature, pH, Dissolved oxygen, NO_2_^-^/NO_3_^-^, NH_3_, PO_4_^3-^, volatile suspended solids, conductivity, and turbidity) between pre-intervention time points (2013-2015) and post-intervention time points (2016-2019). This analysis was conducted using site 76 (immediately downstream of Calumet), site 56 (immediately upstream of Calumet), site 36 (immediately downstream of O’Brien), and site 112 (immediately upstream of O’Brien) (Supplementary figure S3).

Like the fecal coliform data, most of the measured physicochemical variables contained skewed sampling distributions and substantial outliers. Moreover, the physicochemical variables were subject to seasonality, resulting in temporally autocorrelated data. Therefore, we employed another median-based bootstrapping simulation while also using the physicochemical measurements of upstream sites (Site 56 and 112) to account for seasonal variation. For each measured physicochemical variable of interest, differences in the measurements of upstream and downstream sites were calculated for each time point. Then, the median values of such measurements in post-intervention time points (2016-2019) were compared against the median values of such measurements in pre-intervention time points (2013-2016) to form a test statistic. These metrics were compared to a bootstrapped null distribution in which upstream and downstream measurements were drawn from the same sampling pool and the process was repeated to create simulated test statistics under null conditions. The true test statistic was compared against the null distribution in a two-tailed test for significance, and the total p-values were adjusted for multiple comparisons using Benjamini-Hochberg corrections.

## Results

### Microbial community compositions are well defined by sample type

Microbial communities from each sampled medium (water, sediment, sewage, effluent) significantly differed in both alpha and beta diversity indices. Sample types displayed significantly different alpha diversity values as measured by Shannon’s diversity index (Kruskal Wallis test, p < 0.001, Figure 1A). As expected (Sommers et al. 2019, Wang et al. 2012, Jiang et al. 2006, Feng et al. 2009, Payne et al. 2017), sediment samples had the greatest alpha diversity, followed by effluent samples, then followed by water and sewage samples (pairwise Wilcoxon tests with BH corrections; See Figure 1A for statistical groupings.) Notably, there were no significant differences in alpha diversity values between the effluent samples at O’Brien and Calumet, or between the sewage samples at O’Brien and Calumet, although alpha diversity differences between the sewage and effluent types were highly significant (p < 0.001).

**Figure 1.**
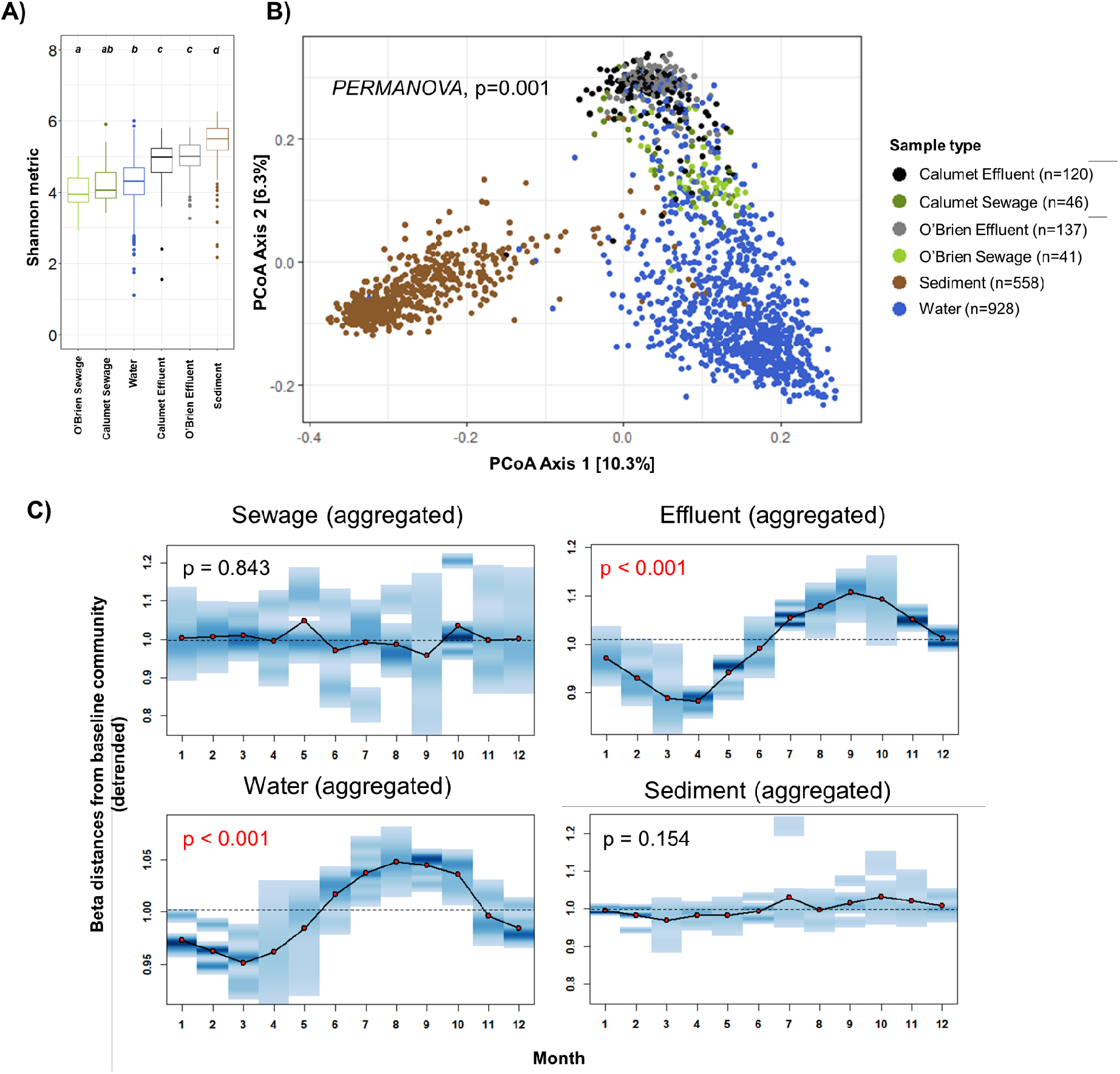
**(A) Alpha diversity of CAWS samples by sample type.** The distribution of Shannon diversity indices for each sample type is consolidated for all seven sampling years (2013-2019). This box and whisker plots demonstrate the quartile range and outliers for each distribution. Statistical groupings are designated by the letters above the boxplots. Significance was assessed using paired Wilcoxon tests with Benjamini-Hochberg multiple testing corrections. **(B) Beta diversity analysis of CAWS samples by sample type.** PCoA plot based on the unweighted UniFrac distance matrix showing clustering patterns of different sample types. **(C) Aggregated seasonal plots of beta distances as a function of month, for each sample type.** The dots connected by solid black lines represent the mean beta distance between each sample community and a fixed baseline community of the same sample type and site. The blue density bands describe the distribution of beta distances to a baseline community, for each particular month. The dashed lines represent the null expectations given that seasonality plays no role in the community. Missing time points were interpolated using spline regression.

Beta diversity analyses using both weighted and unweighted UniFrac distance metrics indicated significant differences in community composition across sample types (weighted UniFrac PERMANOVA p < 0.001, unweighted UniFrac PERMANOVA p < 0.001, Figure 1B). This was supported further by pairwise post-hoc comparisons with Benjamini-Hochberg corrections, which showed significant differences in community composition between all pairwise combinations of sample types (Supplementary Figure S2). Beta dispersion analyses across sample type (calculated as the mean value of unweighted UniFrac distances between all coordinates of individual samples to the coordinate of its respective sample type centroid) indicated that water samples contained the highest levels of spatio-temporal community variability (mean distance to centroid = 0.5759), followed by sediment (mean distance = 0.5499), Calumet sewage (mean distance = 0.5401), Calumet effluent (mean distance = 0.5309), O’Brien effluent (mean distance = 0.5198), and finally O’Brien sewage (mean distance = 0.5123) (See Supplementary Figure S2 for statistical significance).

### Seasonal trends play a dominant role in shaping the CAWS effluent and river community, but not in the sewage and sediment community

The temporal dimension of microbial community dynamics varied across sites and sample media. Traditional ordination-based methods for community distance metrics (*e.g*., PCoA) do not account for the inherent temporal autocorrelation within the data (Martino et al. 2021); therefore, we directly examined the beta distances of each sampled community to a site-specific baseline community over time. We found that several of the site- and sample type-specific time series plots showed a yearly cyclical component (Unweighted Unifrac - Supplemental S2, Weighted UniFrac - Supplemental S3). When aggregating beta distance results across sites, we found evidence for seasonality in effluent and water but not sewage and sediment (Figure 1C). This effect remained clear regardless of the use of unweighted UniFrac distance (effluent samples: Friedman test statistic = 61.00, p-value < 0.001; water samples: Friedman test statistic = 57.13, p-value < 0.001) or weighted UniFrac distance metrics (effluent samples: Friedman test = 57.13, p-value < 0.001; water samples: Friedman test = 47.62, p-value < 0.001). In comparison, we found no evidence for seasonality with sewage (unweighted UniFrac: Friedman test statistic = 6.44, p-value = 0.828; weighted UniFrac: Freidman test statistic = 10.64, p-value = 0.474) or sediment samples (unweighted UniFrac: Friedman test statistic = 15.67, p-value = 0.154; weighted UniFrac: Friedman test statistic = 13.38, p-value = 0.269).

Water and effluent samples contained strong seasonal signals in community composition, therefore we performed compositionally-aware differential abundance analyses (Morton et al. 2019) with both sample types and characterized the specific taxonomic groups driving our observed changes in the community composition across seasons. Differential abundances in ASVs between March and August time points were calculated using Songbird models trained with subsetted data of only water and effluent samples collected at either month (effluent samples n = 63, water samples n = 197). Resultant models contained acceptable goodness-of-fit values for both effluent (pseudo-Q square = 0.161) and water (pseudo-Q square = 0.103) sample types, indicating a significant effect size of month in contributing to model fit. After identifying the top ten ASVs that increased most significantly in March relative to August (henceforth referred to as March-associated ASVs), and top ten ASVs that were differentially abundant in August relative to March (henceforth referred to as August-associated ASVs) (Figure 2A), we examined the log-ratios of March-associated to August-associated ASVs across the entire time series (Figure 2B). We found that such log-ratios indeed followed a gradual and predictable seasonal gradient through time when extrapolated onto the entire time series, providing validation that our identified ASVs follow strong seasonal shifts in relative abundance within their respective sample type.

**Figure 2:**
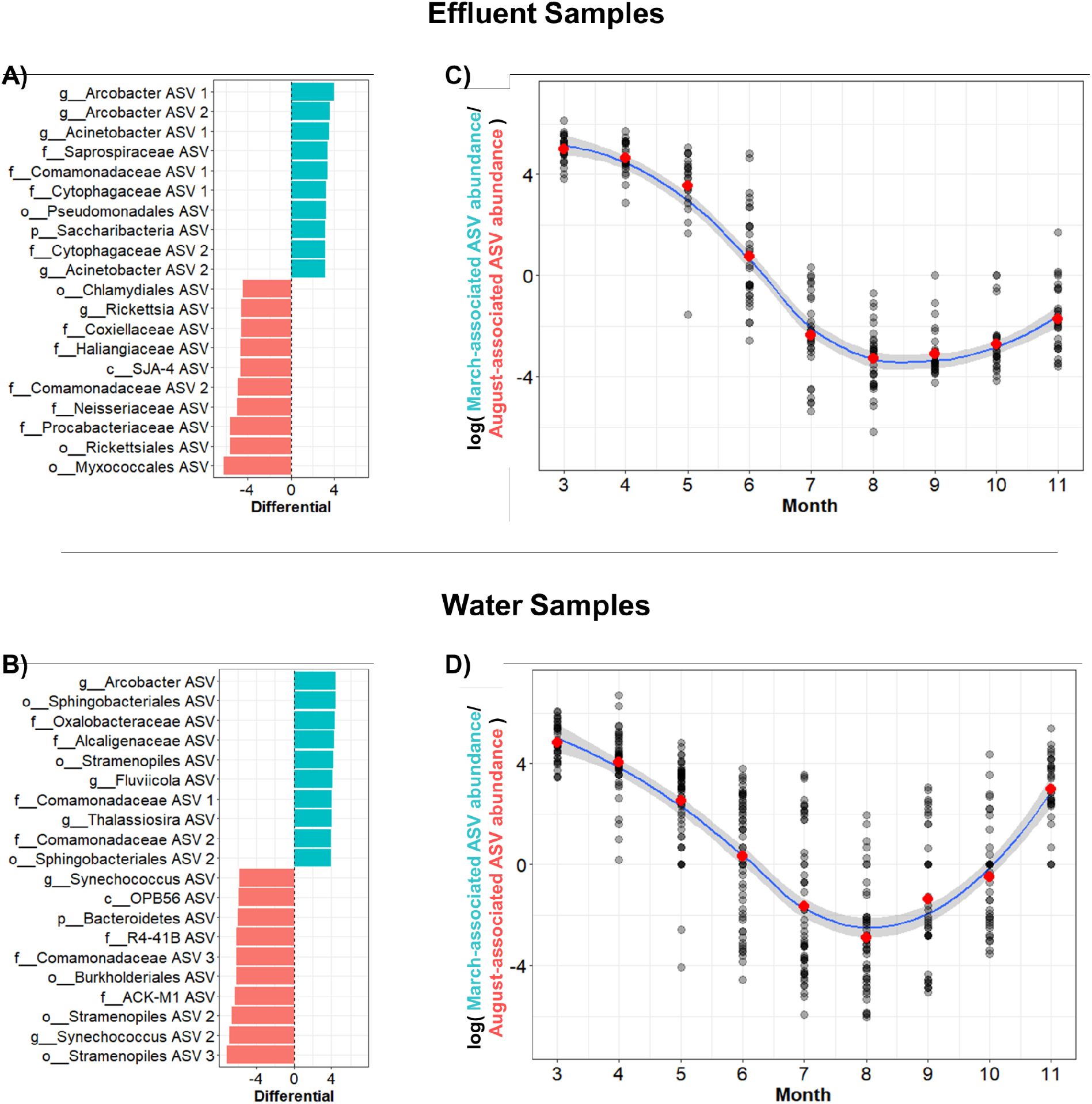
**(A, B) Differential abundance plots of water and effluent samples, comparing samples collected in March to samples collected in August.** Positive differential values indicate the top ten ASVs that were differentially abundant in March samples compared to August, and negative values indicate the top ten ASVs that were differentially abundant in August samples relative to March. **(C, D) Log ratios of March-associated ASVs to August-associated ASVs as a function of month in water and effluent samples.** Red points indicate the mean value of each month. Blue lines are best-fit curves of the data using a local polynomial regression fitting method (loess) with 95% confidence intervals.

### Spatially unique microbial communities experience a continuum of compositional shifts along the river

Time-resolved spatial dynamics of the CAWS microbiome was characterized to examine the impacts of the WRP system upgrade interventions on downstream community dynamics. This targeted analysis was constrained to only include water and sediment samples of sites upstream, immediately downstream, and further downstream of Calumet and O’Brien WRPs, as well the effluent samples from both WRPs (Figure 3A). Significant differences in microbial community structure were quantified between all pairs of sites for both water and sediment samples. (Figure 3B, see Supplemental Table S2 for PERMANOVA results).

**Figure 3:**
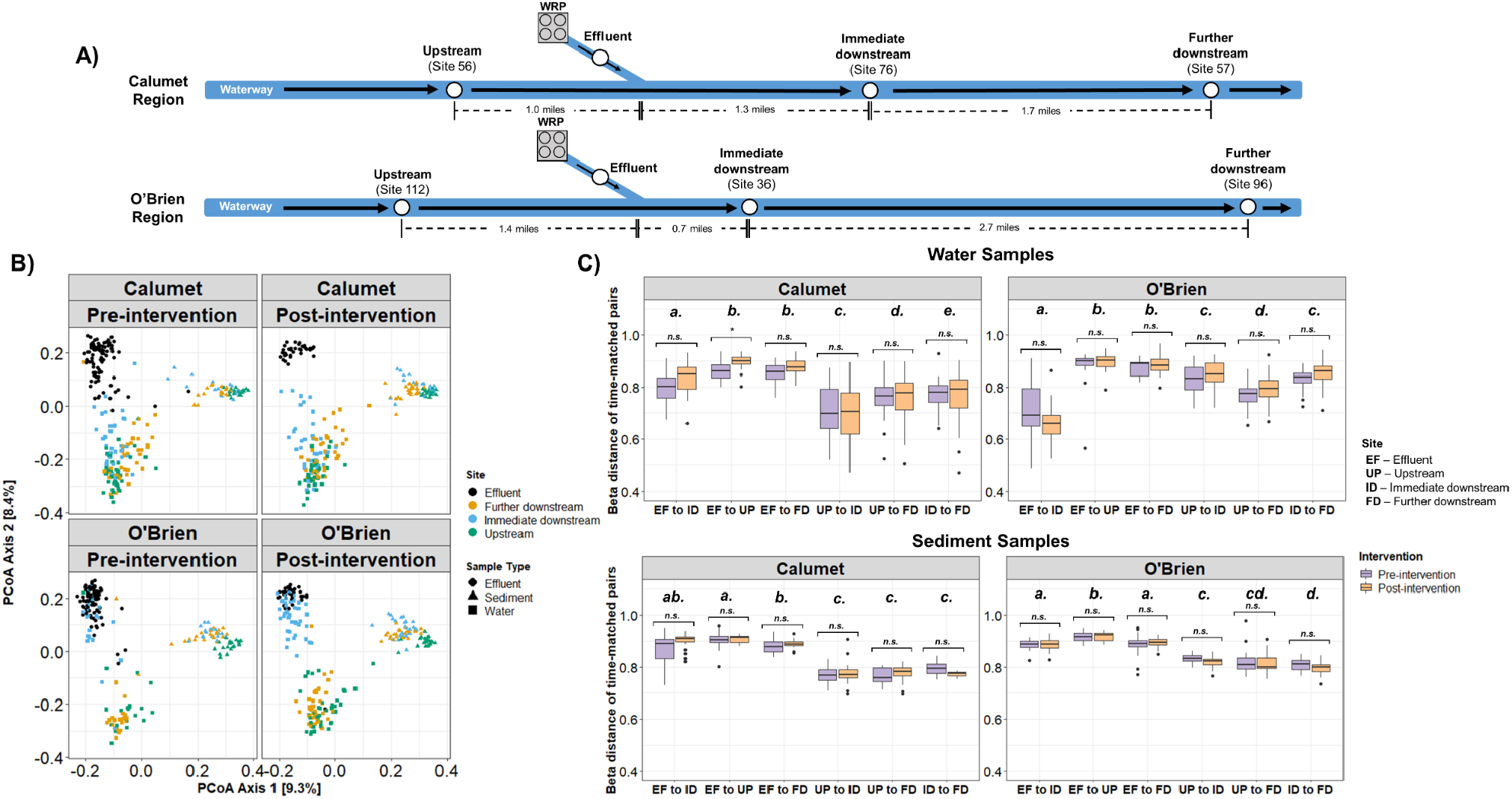
**(A) Simplified map of the Calumet and O’Brien regions of the CAWS.** Subsetted sites used in the following analyses are described in this map. **(B) Unweighted UniFrac PCoA plots of effluent sample, and water and sediment samples from upstream, immediate downstream, and further downstream relative to WRPs.** Plots are separated by region and by intervention period. **(C) Unweighted UniFrac distances between pairs of sites, matched by time point.** Statistical comparisons between pre- and post-intervention distances within each pair are denoted using brackets, with n.s. indicating non-significance (p>0.05). Statistical groupings of comparisons across pairs are designated by the letters above the boxplots. Significance was assessed using t-tests with Benjamini-Hochberg multiple-testing corrections.

Examinations of time-resolved beta distances between WRP-adjacent sites showed that community similarity measures were often consistent with the spatial orientation of site locations to one another and to the WRP outfalls, particularly with water samples (Figure 3C). In the O’Brien region, water communities collected from the site immediately downstream of the WRP were significantly more similar to the effluent communities than water samples upstream or further downstream. Further downstream from the O’Brien WRP, water communities became less similar to effluent samples and increased in resemblance to the upstream river community prior to WRP effluent outflow. We found similar trends in the Calumet region as well, with water communities immediately downstream of the WRP being the most compositionally similar to effluent compared to the other river sites. However, unlike the case in the O’Brien region, the community distances of the immediate downstream site to effluent were significantly larger compared to its distances to the other river sites, likely owing to the fact that the immediate downstream site at the Calumet region is twice as far from the WRP outfall than at the O’Brien region.

Within sediment communities, we found that each site also contained compositionally distinct microbial communities whose differences remained robust through time (Supplementary Table S2, Figure 3C). Additionally, we note that unlike in the water communities, sediment community similarities did not follow a spatial orientation consistent with site location; microbial communities from all sites had mostly the same beta diversity distance to the effluent, for both the Calumet and O’Brien regions (Figure 3C). Similar results were found when measuring beta distances with weighted UniFrac distances (Supplemental figure S5).

We found minimal evidence that the WRP upgrades caused any significant changes in site-specific community compositions in either water or sediment. Comparisons of time-resolved beta distances between pre and post-intervention time points found a statistically significant change in distance for only one site pair: effluent to upstream river communities at the Calumet region (Figure 3C). Re-analysis with weighted UniFrac distances also did not identify meaningful community changes as a result of the intervention (Supplemental figure S5). Unsurprisingly, differential abundance models attempting to identify key microbial ASVs that significantly changed in abundance between pre-intervention to post-intervention time points resulted in poor model fits (pseudo Q square scores < 0.05), indicating low effect sizes.

### The WRP upgrades are effective in reducing specific populations of coliform-forming bacteria in the river, yet observable effects quickly diminish downstream

The spatio-temporal trends of fecal coliform counts cultured from water samples were analyzed. These counts are used as indicators of potential fecal contamination and as metrics for overall water quality for recreational and drinking use (McLain and Williams 2012). Throughout the study period fecal coliform concentrations (measured as the number of colony forming units per 100 mL of sample, or CFU/100mL) were quantified from effluent and water samples. In the effluent samples, bootstrap hypothesis tests showed that median values of fecal coliform concentrations were significantly and substantially reduced following water treatment upgrades at both O’Brien (absolute effect size = 10,940 CFU/100mL, p < 0.001) and Calumet (absolute effect size = 8280 CFU/100mL, p < 0.001) sites. All pre-upgrade effluent samples contained fecal coliform concentrations that considerably exceeded the recreational water quality standard of 400 CFU/100mL (USEPA 1986) while all post-upgrade effluent samples contained concentrations below this value. When examining coliform concentrations of river sites immediately downstream of Calumet and O’Brien WRPs, bootstrapping tests also found a significant reduction in coliform concentrations post upgrade; however, the effect size was weaker relative to the effluent samples and the signal contained higher noise (Calumet effect size = 1940 CFU/100mL, p < 0.001, O’Brien effect size = 5690 CFU/100mL, p = 0.004). Finally, examinations of coliform concentrations in sites further down the river contained no significant differences in concentrations between pre- and post-upgrade time points (p>0.05), with levels fluctuating continually between below to above the EPA standard.

### WRP upgrades did not significantly alter the physio-chemical environment of the downstream CAWS

Microbial community composition and dynamics are influenced by physicochemical parameters such as temperature, oxygen, nutrients, and pH. We tested whether WRP upgrades caused significant changes in key physicochemical water measurements in the downstream river sites by calculating the differences in physicochemical conditions at upstream and downstream sites relative to WRPs at each time point. We then tested for significant changes in such upstream-downstream differences between pre-intervention time points and post-intervention time points, using a median-based bootstrap hypothesis test (Table 1). Overall, we found no statistical support that the WRP upgrades caused significant shifts in physicochemical differences between upstream and downstream sites. In both the Calumet and O’Brien regions, we found no significant evidence that the intervention influenced upstream-downstream differences in any of the measured physicochemical variables. These comparisons of physicochemical parameters suggest the WRP upgrades played a relatively minimal role in affecting the physical and chemical conditions of downstream sites, particularly when compared to the natural seasonal variability captured at upstream sites (Figure S3).

**Table 1:** Results of median-based bootstrap hypothesis tests measuring for significant changes in physicochemical parameters between pre- and post-intervention time points, at Calumet and O’Brien WRP sites.

|  | Calumet region |  |  | O'Brien Region |  |  |
| --- | --- | --- | --- | --- | --- | --- |
|  | Bootstrap test statistic | p-value | adjusted p-value | Bootstrap test statistic | p-value | adjusted p-value |
| Chlorophyll (ug/L) | 1.6 | 0.433 | 1.000 | -0.6 | 0.807 | 1.000 |
| Dissolved oxygen (mg/L) | 0.25 | 0.514 | 1.000 | 0.215 | 0.623 | 1.000 |
| NO<sub>2</sub>/NO<sub>3</sub> (mg/L) | 0.1745 | 0.486 | 1.000 | 0.52 | 0.565 | 1.000 |
| NH<sub>3</sub> (mg/L) | -0.165 | 0.009 | 0.101 | -0.19 | 0.614 | 1.000 |
| Total phosphorus (mg/L) | -0.21 | 0.514 | 1.000 | 0.573 | 0.057 | 0.627 |
| Volatile suspended solids (mg/L) | 0 | 1.000 | 1.000 | 0 | 1.000 | 1.000 |
| Total organic carbon (mg/L) | -0.2 | 0.284 | 1.000 | 0.05 | 0.762 | 1.000 |
| Conductivity (mS) | 0.0195 | 0.484 | 1.000 | -0.0355 | 0.560 | 1.000 |
| Turbidity (NTU) | 3.475 | 0.022 | 0.246 | -3.075 | 0.255 | 1.000 |
| pH | 0.145 | 0.051 | 0.557 | -0.075 | 0.479 | 1.000 |
| Temperature (°C) | 0.65 | 0.132 | 1.000 | 0.4 | 0.502 | 1.000 |

## Discussion

This study, which spanned 7 years from 2013 to 2019, aimed to characterize the spatio-temporal dynamics of microbial communities associated with the CAWS using 16S rRNA gene amplicon sequencing of samples collected from sediment, water, treated effluent, and raw sewage, along with fecal coliform counts and physicochemical measurements. Additionally, we investigated the impact of MWRD water quality improvement efforts (i.e. the WRP disinfection system upgrades and the implementation of the TARP system to capture CSOs) on the community dynamics of the CAWS microbiome.

As expected, compositional analysis of microbial communities demonstrated distinct distribution patterns across environmental media (river water, sediment, effluent, sewage), with significant differences between sample types for both alpha and beta diversity. The microbiomes of each sample type were compositionally unique, with 14.8% of the variance of community composition being explainable by sample type alone when using unweighted UniFrac distances (and 30.5% when using abundance-weighted UniFrac distances). Such results are in line with expectations that, while all sampled environments are interconnected within the urban river ecosystem, each environmental medium still harbors a distinct microbial community shaped by unique selection forces. As a noteworthy example, the microbial communities found in water samples contained relatively low alpha diversity values but the highest levels of spatio-temporal variability in community composition, consistent with expectations of high community turnover and shifting selection pressures typically found in aquatic environments (Payne et al. 2017, Savio et al. 2015). Sediment microbiomes were found to be most different from the other sample types (Figure 1); as river water, sewage, and effluent all share the similarity of being aquatic environmental media, this reinforces the role of substrate type as a primary driver for community differences (Lozupone and Knight 2007, Thompson et al. 2017).

Effluent samples and sewage samples significantly differed in both diversity-related and composition-related characteristics at both WRP sites, demonstrating the sizable impact of wastewater processing in affecting the microbial communities present in wastewater. At wastewater reclamation plants, sewage is initially turned into activated sludge, an enrichment culture of microorganisms that can remove biological oxygen demand and total suspended solids from wastewater (Nascimento et al. 2018, VandeWalle et al. 2012), then processes such as aeration and final disinfection steps can lead to enrichment of sewage treatment-associated microorganisms and overall changes in community composition (Newton et al. 2022). Interestingly, we also note that samples from the two WRP sites were very similar to one another when examining both effluent and sewage sample types. For example, alpha diversity metrics were statistically indistinguishable between the two sites (Figure 1) and pairwise PERMANOVAs found significant differences yet relatively low effect sizes when comparing site pairs of the same sample type (Table S1); this indicates that both WRP regions receive similar microbial communities as sewage input, and that both treatment procedures tend to produce microbially similar effluent outputs despite using different disinfection strategies.

Spatiotemporal examinations of the CAWS community revealed that the wastewater treatment and CSO capture upgrades did not have dramatic effects on the overall community of the downstream river; rather, the effects were nuanced and population-specific. Proximity of water sampling location to effluent outfall exerted a clear influence on microbial community structure; for both Calumet and O’Brien, microbial communities in water immediately downstream of the outfall were consistently and significantly more similar to effluent than communities from other water samples. This effect was most pronounced for O’Brien, where the “immediate downstream” site was only 0.7 miles downstream of the outfall, compared to the closest downstream site to the Calumet outfall which was 1.3 miles. Indeed, for the O’Brien system, the immediate downstream water samples contained microbial communities that resembled effluent more than they resembled upstream or downstream water from the river, while the site immediately downstream of the Calumet outfall was influenced by effluent to a lesser degree and most closely resembled the site upstream. Despite the wastewater treatment system upgrades at both WRPs in 2016, beta distances between downstream water and effluent microbial communities remained consistent before and after the upgrades, suggesting that these upgrades did not exert a strong influence on changing the relative compositions of downstream microbial communities.

Measurements of fecal coliform concentration, a traditional standard of freshwater quality (Gronewold et al. 2008, Meays et al. 2004, USEPA 1984), demonstrated the clear success of the intervention in reducing the absolute abundance of fecal-associated bacterial taxa from effluent samples. This was also found to be the case with downstream river sites; yet, similarly to the beta distance analysis, effectiveness tended to diminish with respect to the samples’ distance from the WRP sites. In fact, sites greater than 3 miles away from either WRP no longer had any detectable differences in fecal coliform concentrations between pre- and post-intervention time points, remaining at a level comparable to pre-intervention timepoints and frequently exceeding the EPA recreational water threshold. Such results may indicate the presence of o ther sources of fecal coliform-associated bacteria to the waterway besides the WRP effluent and pre-TARP CSO events. We additionally note that this described spatio-temporal pattern was nearly identical between the Calumet and O’Brien regions, demonstrating the comparable effectiveness of both UV-based and chlorination-based disinfection protocols in reducing fecal coliform concentrations from treated wastewater.

In contrast to fecal coliform results, we found that the overall dynamics of CAWS microbial communities appeared robust and consistent over the course of the study. Neither PERMANOVA nor time-resolved beta distance analyses using site- and sample-specific comparisons revealed any significant impact of the system upgrades in the compositional community dynamics of downstream river sites. Instead, the predominant driver of microbial community composition was a consistent yearly cyclical trend, a trend which we attribute to seasonal effects driving the ecology of the CAWS microbiome. This was particularly the case with effluent and water samples, while sediment and sewage communities remained consistent throughout each year. Evidence for seasonal effects have been observed in various aquatic-based microbiomes, most prominently in marine systems (Bunse and Pinhassi 2017, Gilbert et al. 2012) but additionally in freshwater river systems (Staley et al. 2015, Zhang et al. 2011) and in the activated sludge of wastewater systems (Johnson et al. 2020, Pesces et al. 2022, Ju et al. 2014). In our study we found that such seasonal effects overpowered any potential effect of the intervention in shifting community dynamics of downstream river sites. Similar examples have been noted in aquaculture systems (Marmen et al. 2021, Zeng et al. 2019), indicating the pervasiveness of seasonality as a major contributor to microbial community structure. We also found that there was relatively minimal overlap in the specific ASVs most affected by seasonality when comparing between effluent and water samples, suggesting that microbial populations most significantly affected by seasonality are community and environment specific.

Notably, we found that sewage and sediment did not appear to be affected by seasonal trends in our study (although there were a few exceptions of site-specific sediment communities that tested significantly for seasonality). Of particular note is our finding that the sewage communities in both O’Brien and Calumet WRPs did not appear to contain a discernible relationship to time of year, despite the highly consistent seasonal signal found in the effluent. We also note that such results contrast with other work examining the sewage microbiome in other locales (LaMartina et al. 2021). As engineered microbiomes that are introduced to activated sludge during wastewater treatment have demonstrated seasonal qualities (Johnson et al. 2020, Pesces et al. 2022, Ju et al. 2013), we speculate that the wastewater treatment process in our system may introduce a seasonal component to the microbial communities as they transition from untreated sewage to treated effluent, although confirmation would require a more nuanced study. As for the sediment, communities were found to be relatively stable through time and instead were most definable by site, suggesting that despite their close proximity to water, their dynamics are distinct from those of water.

River systems are unique environments for examining microbial community dynamics, in that they offer spatial complexity that is integrated with temporal patterns. In this seven-year study, we characterized the microbial community of the CAWS across a broad spatio-temporal gradient and described its response to a prominent environmental change. Our results from this seven-year long microbiome study are nuanced; on one hand, we provide evidence that wastewater management improvement efforts such as implementation of large scale disinfection technologies and combined sewer overflow capture systems can lead to significant improvement in water quality, as indexed by reductions in population sizes of known indicators of fecal contamination. On the other hand, our analyses indicate that such water quality improvement measures do not appear to greatly shift the structure of existing microbial community dynamics in the waterway overall, or the physicochemical environment in which the microbial communities exist. Our results demonstrate the robustness of community dynamics in the system despite an interventional disturbance that significantly reduced the prevalence of fecal coliform-associated bacteria immediately downstream of the WRPs.

**Figure 4:**
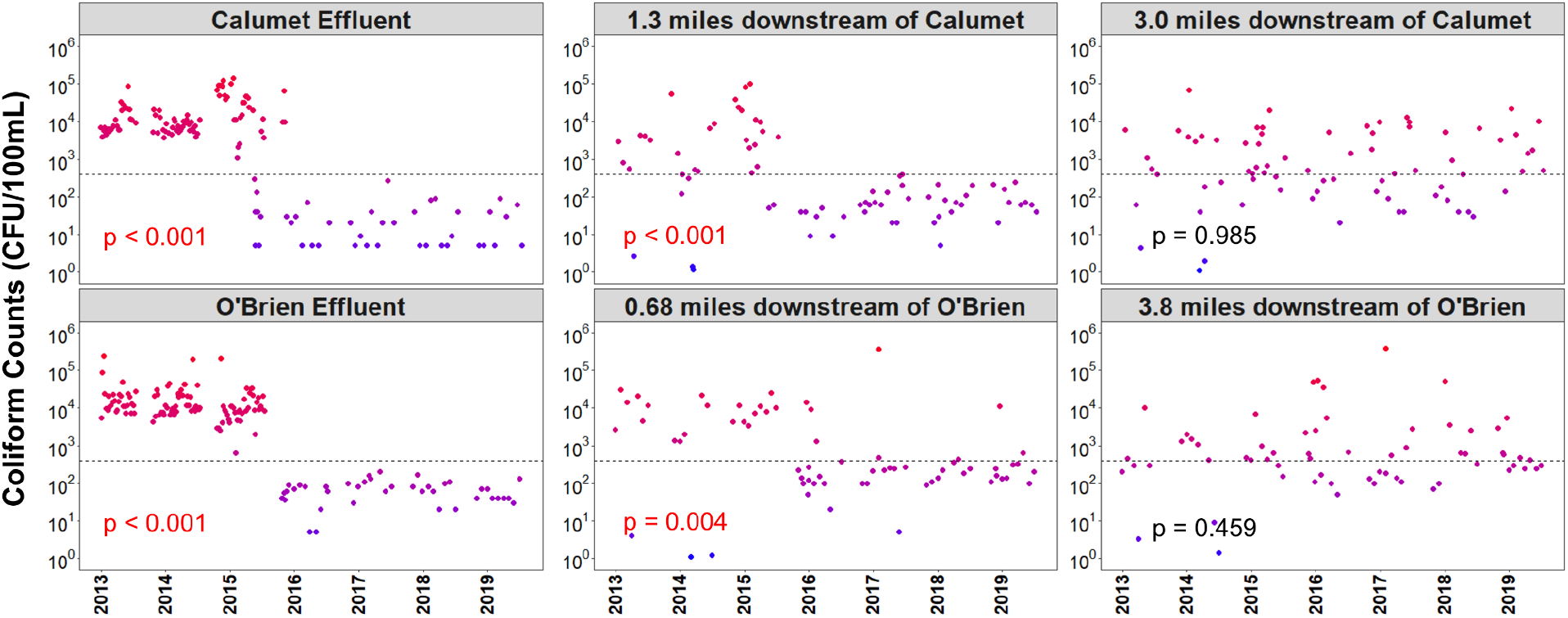
Log-transformed fecal coliform concentrations as a function of time in effluent and downstream river samples of Calumet and O’Brien WRPs. Plots on the left represent effluent samples directly from WRPs, plots in the center represent samples of river sites immediately downstream from WRPs, and plots on the right represent samples of river sites further downstream of the WRPs. The dotted line represents 400 CFUs per 100mL, a concentration set as the EPA standard for recreational waters. The p-values represent results of bootstrapping simulations testing for differences in median values of coliform concentrations between pre-intervention time points (before 2016) and post-intervention time points (after 2016).

## Glossary/Key Definitions

CAWS: The Chicago Area Waterway System.
WRP: Wastewater reclamation plant. There are two WRPs in the system, named O’Brien and Calumet.
Sewage: Incoming untreated wastewater entering WRPs.
Effluent: Outgoing treated discharge leaving WRPs.
CSO: Combined Sewer Overflow event, leading to discharges of untreated sewage directly into the CAWS.
TARP: Tunnel and Reservoir Plan, aimed to reduce the number of CSO events.
Intervention: The implementation of disinfection upgrades at WRPs as well as TARP upgrades to the CAWS system in 2016.

## Data Availability

The dataset supporting the conclusions of this article is available in the European Bioinformatics Institute repository, [ERP136279]. Additionally, sequencing data and processed tables and taxonomy assignments are available through QIITA under study ID14446.

## Author contributions

JAG and CN conceived of the study. MG collected processed samples. MD and AS conducted bioinformatic processing of resultant data. SMK conducted formal analysis and visualization. SMA, HLL, CM, RK, and JAG supervised and provided feedback on formal analysis and visualization. SMK wrote the manuscript, and all authors edited and approved the final manuscript.

## Acknowledgements

This work would not have been possible without the support of the Chicago Metropolitan Water Reclamation District.

